# Identification of a non-canonical function of prefoldin subunit 5 in proteasome assembly

**DOI:** 10.1101/2024.04.22.590501

**Authors:** Somayeh Shahmoradi Ghahe, Krzysztof Drabikowski, Ulrike Topf

## Abstract

The prefoldin complex is a heterohexameric, evolutionarily conserved co-chaperone that assists in the folding of polypeptides downstream of the protein translation machinery. Loss of prefoldin function leads to impaired solubility of cellular proteins. The degradation of proteins by the proteasome is an integral part of protein homeostasis. Failure of regulated protein degradation can lead to the accumulation of misfolded and defective proteins. We show that prefoldin subunit 5 is required for proteasome activity by contributing to the assembly of the 26S proteasome. In particular, we found that the absence of prefoldin subunit 5 impairs the formation of the Rpt ring subcomplex of the proteasome. Concomitant deletion of *PFD5* and *HSM3*, a chaperone for assembly of the ATPase subunits comprising the Rpt ring, exacerbates this effect, suggesting a synergistic relationship between the two factors in proteasome assembly. Thus, our findings reveal a regulatory mechanism wherein prefoldin subunit 5 plays a crucial role in maintaining proteasome integrity, thereby influencing the degradation of proteins.

## Introduction

Prefoldin, a co-chaperone conserved across eukaryotes and archaea, facilitates the co-translational folding of cytoskeletal actin and tubulin by transferring their nascent chains to the TRiC/CCT complex (Vainberg et al., 1998). This interaction enhances the efficiency of folding reactions (Geissler et al., 1998; Gestaut et al., 2019) and is independent of ATP hydrolysis. Prefoldin not only binds and stabilizes nascent polypeptides but also interacts with various protein substrates involved in cellular pathways such as chromatin remodeling, transcription regulation, RNA splicing (Costanzo et al., 2016; Payan-Bravo et al., 2021), protein turnover, and cancer progression (Herranz-Montoya et al., 2021; Tahmaz et al., 2021). Structurally, eukaryotic prefoldin comprises six distinct subunits: two α-like (Pfd3/Gim2/Pac10 and Pfd5/Gim5) and four β-like (Pfd1/Gim6, Pfd2/Gim4, Pfd4/Gim3, and Pfd6/Gim1) subunits (Leroux et al., 1999; Martin-Benito et al., 2002; Vainberg et al., 1998). Like its archaeal homolog, eukaryotic prefoldin adopts a jellyfish-like structure, with subunits resembling six tentacles consisting of two long coiled coils extending from a base of β barrels. The distal ends of coiled coils contain unique charged residues that serve as interaction sites for binding and stabilizing substrates (Martin-Benito et al., 2002). Each prefoldin subunit interacts with a specific group of substrates, potentially influencing particular cellular pathways (Simons et al., 2004). The prefoldin complex participates in various intracellular mechanisms, and deficiencies in it have been associated with pathological conditions, including neurodegenerative diseases resulting from the intracellular accumulation of toxic aggregates (Liang et al., 2020; Tashiro et al., 2013).

Unfolded and misfolded proteins, unable to refold, may acquire new anomalous functions through aggregate formation. Failure to clear these aggregates compromises proteome integrity and drives pathological conditions (Mathieu et al., 2020; Vaquer-Alicea & Diamond, 2019). In eukaryotic cells, the ubiquitin-proteasome system (UPS) and selective autophagy manage the removal of aberrant proteins and toxic aggregates (Varshavsky, 2017). The UPS primarily degrades soluble misfolded and redundant proteins tagged by ubiquitin conjugation (Bard et al., 2018; Meyer-Schwesinger, 2019). At the culmination of this system lies the 26S proteasome, responsible for recognizing and degrading ubiquitinated substrates (Dong et al., 2019). Several studies have associated deficiencies in the prefoldin complex with the propagation of protein aggregates (Abe et al., 2013; Glickman et al., 1998; Sorgjerd et al., 2013; Takano et al., 2014; Tashiro et al., 2013). However, the mechanistic understanding of prefoldin’s role during protein degradation remained unclear.

In this study, we used the yeast *Saccharomyces cerevisiae* to decipher the role of the prefoldin complex in the maintenance of cellular proteostasis at physiological and heat stress conditions. In particular, we found that loss of prefoldin subunit 5 resulted in reduced assembly and activity of the 26S proteasome, despite overproduction of proteasome subunits. Native gel analysis revealed a defect in the assembly of the base of the 19S regulatory particle. Consistent with this, our data showed that *PFD5* genetically interacts with *HSM3*, a 19S regulatory particle assembly chaperone, and that a double deletion increases the assembly defect. Thus, our analysis provides evidence for the involvement of Pfd5 in proteasome assembly and proposes a mechanism to explain the accumulation of ubiquitinated protein species upon limited function of the prefoldin complex.

## Results

### Prefoldin subunit 5 is necessary for 26S proteasome activity

To explore the role of prefoldin complex in preventing protein aggregation, we decided to analyse the activity and assembly of the proteasome in prefoldin mutant strains. To exclude additive effects caused by cellular stress conditions, we performed the analysis at physiological temperature of 25°C. Native extracts of all single prefoldin subunit deletions and wild-type cells were used to analyse proteasome proteolytic activity by hydrolysis of a fluorogenic substrate (Figure 1A). The deletion of prefoldin subunit 5 was the only single deletion among all other prefoldin subunits that showed decreased activity of the 26S proteasome, comprising one core particle (CP) and two regulatory particles (RP), referred to as doubly-capped (RP2CP) proteasome (Figure 1A, left panel). At the same time, Δ*pfd5* showed increased activity and levels of core particle (CP) and Blm10-CP, which is a complex of CP and its positive regulator, Blm10 (Dange et al., 2011; Schmidt et al., 2005) (Fig. 1A right panel, Fig. 1B). To prove that the absence of Pfd5 is the primary cause of the defective 26S proteasome, we transformed Δ*pfd5* cells with a construct expressing *PFD5* under the control of its native promoter or empty vector (EV) as a control. We assessed the growth rate of the transformants using two independent transformants of Δ*pfd5* cells expressing *PFD5* and two transformants with EV as well as wild-type cells transformed with EV. We observed that expression of *PFD5* could rescue the growth defect phenotype of *PFD5* deletion strains (Supplementary Figure 1A).

**Fig. 1.**
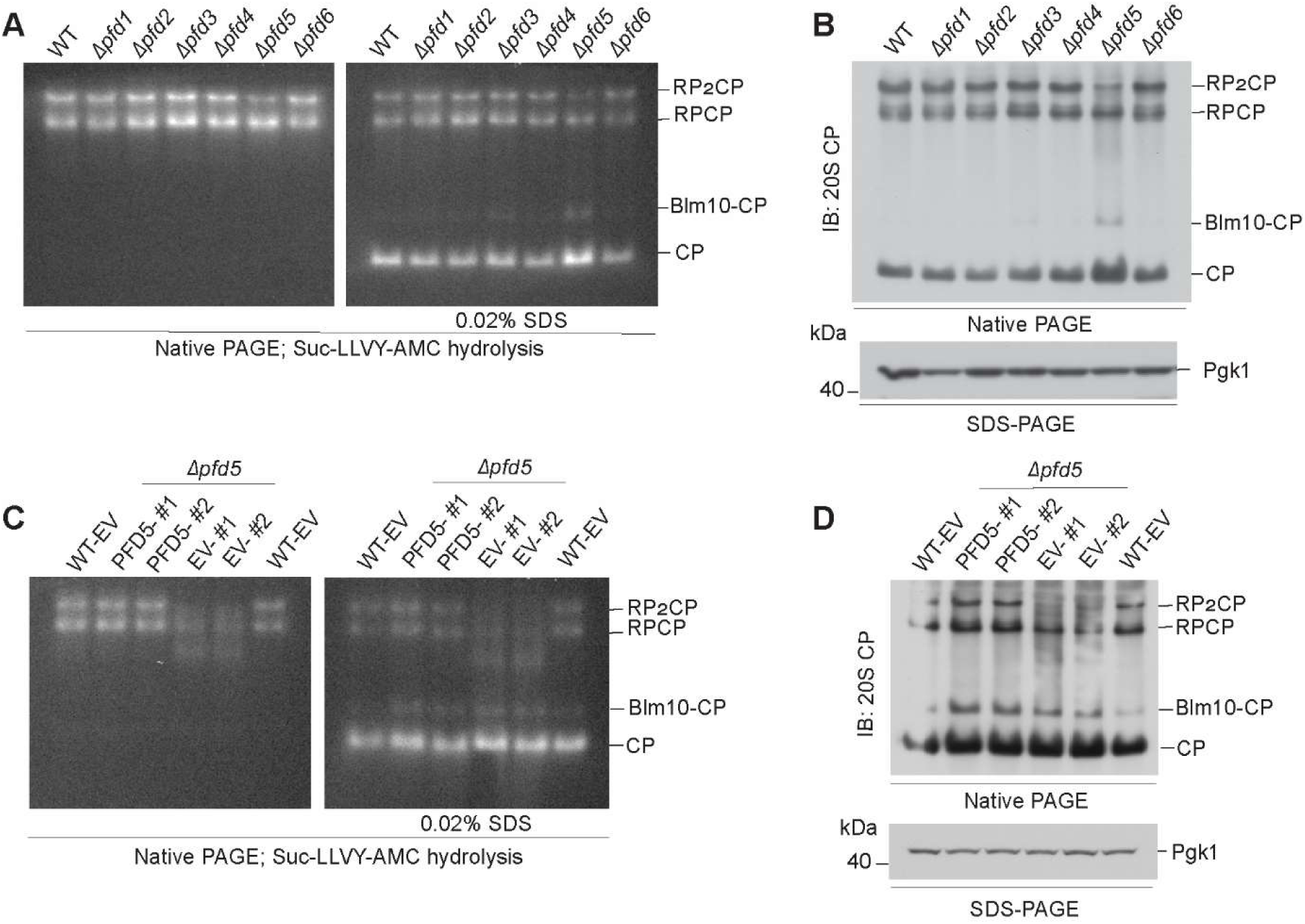
Pfd5 supports 26S proteasome activity. **A, B.** Wild type and *PFD* deletion strains (Δ*pfd1-6*) were grown on minimal medium containing glycerol at 25°C. Equal concentration of cell lysates were resolved by native-PAGE to assess the proteasome proteolytic activity by Suc-LLVY-AMC hydrolysis (A) and by immunoblotting (B). **C, D**. The Δ*pfd5* cells were transformed with a vector expressing *PFD5* under its native promoter or empty vector (EV). Wild-type cells were also transformed with EV. Transformed cells were grown on selective medium containing glycerol at 25°C to logarithmic growth phase. Equal concentrations of cell lysates were separated by native-PAGE to assess the proteasome proteolytic activity by Suc-LLVY-AMC hydrolysis (C) and immunoblotting (D). WT, wild type. EV, empty vector. #1 and #2, two different yeast transformants.

Next, we tested the effect of *PFD5* expression on the activity and assembly of proteasome complexes by in-gel peptidase assay and immunoblotting. EV-transformed Δ*pfd5* cells showed mainly CP peptidase activity and very low levels of single-capped proteasome (RPCP) activity, as well as some intermediates that migrated faster than RPCP, while RP2CP activity was undetectable (Fig. 1C). Complementation with *PFD5* expression could partially restore the peptidase activity of RP2CP, reduced the levels of assembly intermediates and formed a distinct RPCP band (Figure 1C). These observations were confirmed by immunoblotting of 20S CP (Figure 1D), which clearly showed that complementation of Δ*pfd5* cells with plasmid-born *PFD5* could rescue the assembly of RP2CP complexes. Immunoblotting of Rpt1, Rpt5, Rpn5 and Rpn10 (subunits of RP) showed that in Δ*pfd5* -EV cells, the levels of RPCP were insufficient and limited the formation of the RP2CP proteasome holoenzyme (Supplementary Figure 1B). In contrast, expression of *PFD5* in Δ*pfd5* cells rescued the formation of RPCP and assembly of RP2CP proteasome holoenzyme to levels similar to those in wild-type cells (Supplementary Figure 1B). Thus, our data showed that prefoldin subunit 5 is necessary for the sufficient assembly of active 26S proteasome.

### Pfd5-dependent defect of the proteasome is largely independent of ROS levels

We have recently shown that loss of *PFD5* affects the growth of cells on respiratory medium and leads to impaired mitochondrial morphology (Tahmaz et al., 2023). An increase in oxidative stress has been shown to lead to disassembly of 26S proteasome (Reincheckel et al., 1998; Reinheckel et al., 2000). To analyse whether increased production of reactive oxygen species (ROS) due to the mitochondrial defect explains the 26S proteasome assembly defect in Δ*pfd5* cells, we assessed the extent of oxidative stress in prefoldin subunit deletions. All prefoldin subunit deletions showed only a trend towards increased ROS levels compared to wild-type cells (Figure 2A). However, there was no detectable effect on protein carbonylation, a marker for irreversible protein damage caused by oxidative stress (Fig. 2B). We reasoned that if oxidative stress in Δ*pfd5* cells affects proteasome assembly, ROS scavenging might prevent the defect. We treated wild-type and deletion cells with the ROS scavenger N-acetylcysteine (NAC), but could not detect a recovery of proteasome activity compared to untreated cells (Fig. 2C). Furthermore, NAC treatment did not restore the assembly of the 26S proteasome (RP2CP; Fig. 2D). We concluded that oxidative stress is not the primary cause of the proteasome assembly defect in prefoldin subunit 5 deletion strain.

**Fig. 2.**
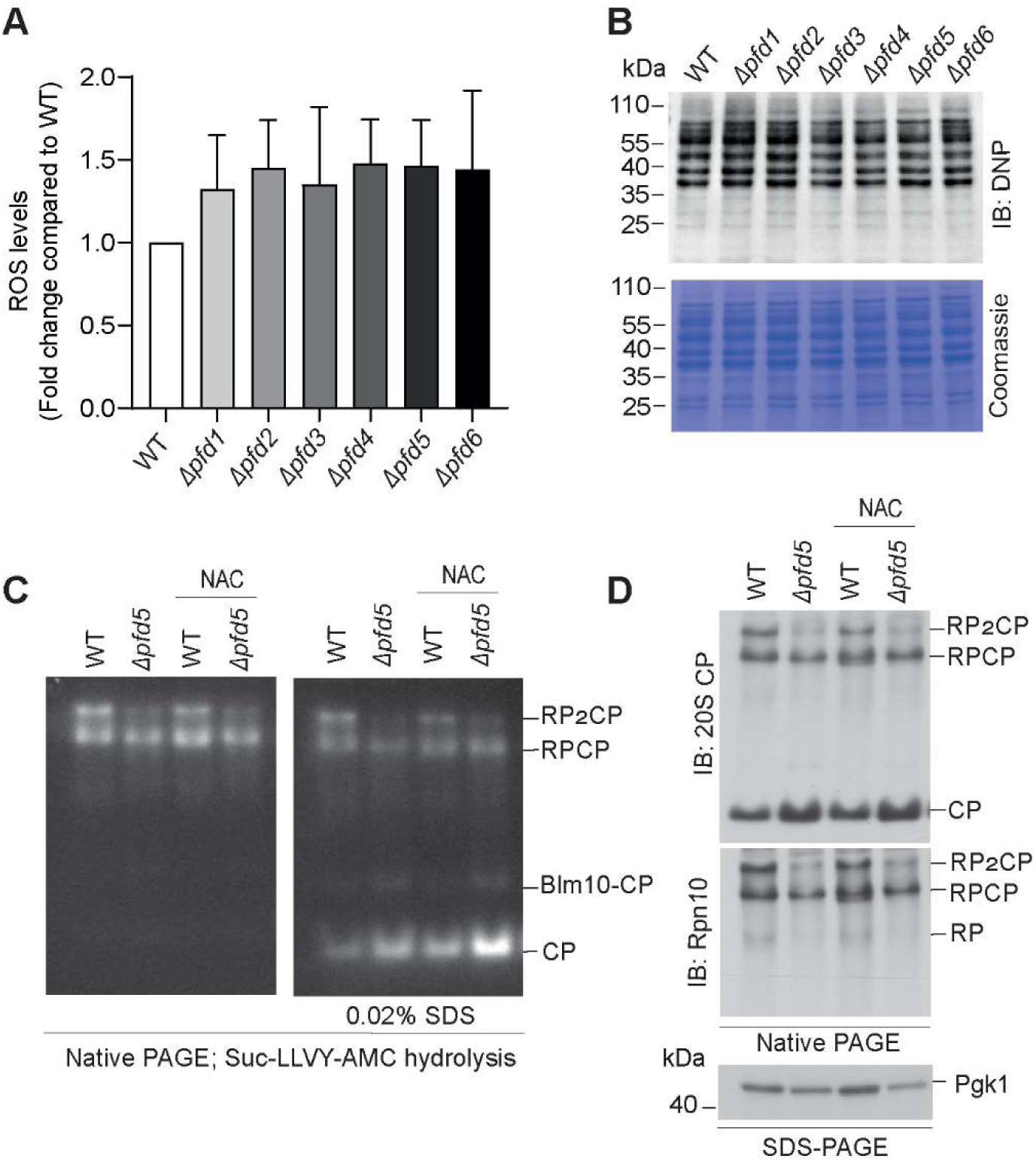
Proteasome assembly defect in Δ*pfd5* is not caused by increased levels of reactive oxygen species. **A.** Levels of reactive oxygen species were determined by incubation of yeast cells with dihydroethedium. The data are represented as mean ± SEM, *n* =5. **B.** Carbonylation of proteins was determined by derivatisation of total yeast cell lysates with DNPH. Equal concentrations of cell lysates were separated by SDS-PAGE and membrane was probed with anti-DNP antibody. Coomassie staining was used as loading control. **C, D.** Yeast cells grown in logarithmic phase were treated with 100 mM N-acetylcysteine (NAC) for four hours. Equal concentration of cell lysates were resolved by native-PAGE to assess the proteasome proteolytic activity by Suc-LLVY-AMC hydrolysis (C) and by immunoblotting (D).

### Deletion of *PFD3* and *PFD5* upregulates proteasome subunits

We tested whether the lack of sufficient amounts of proteasome subunits could be a possible cause of the proteasome defect in Δ*pfd5* cells. First, we analysed the transcript levels of selected proteasome subunits by quantitative RT-PCR. We examined the expression levels of several RP subunits, including *RPT1*, *RPT5* and *RPN1*, subunits of the base subcomplex, as well as *RPN5* of the lid complex and *RPN10*, the proteasome polyubiquitin receptor at the interface of lid and base. The data showed a higher tendency of Δ*pfd3* and Δ*pfd5* strains to upregulate genes encoding proteasome subunits compared to the wild type, with *RPN1*, *RPN10* and *RPT1* being significantly increased in Δ*pfd5* (Figure 3A).

**Fig. 3.**
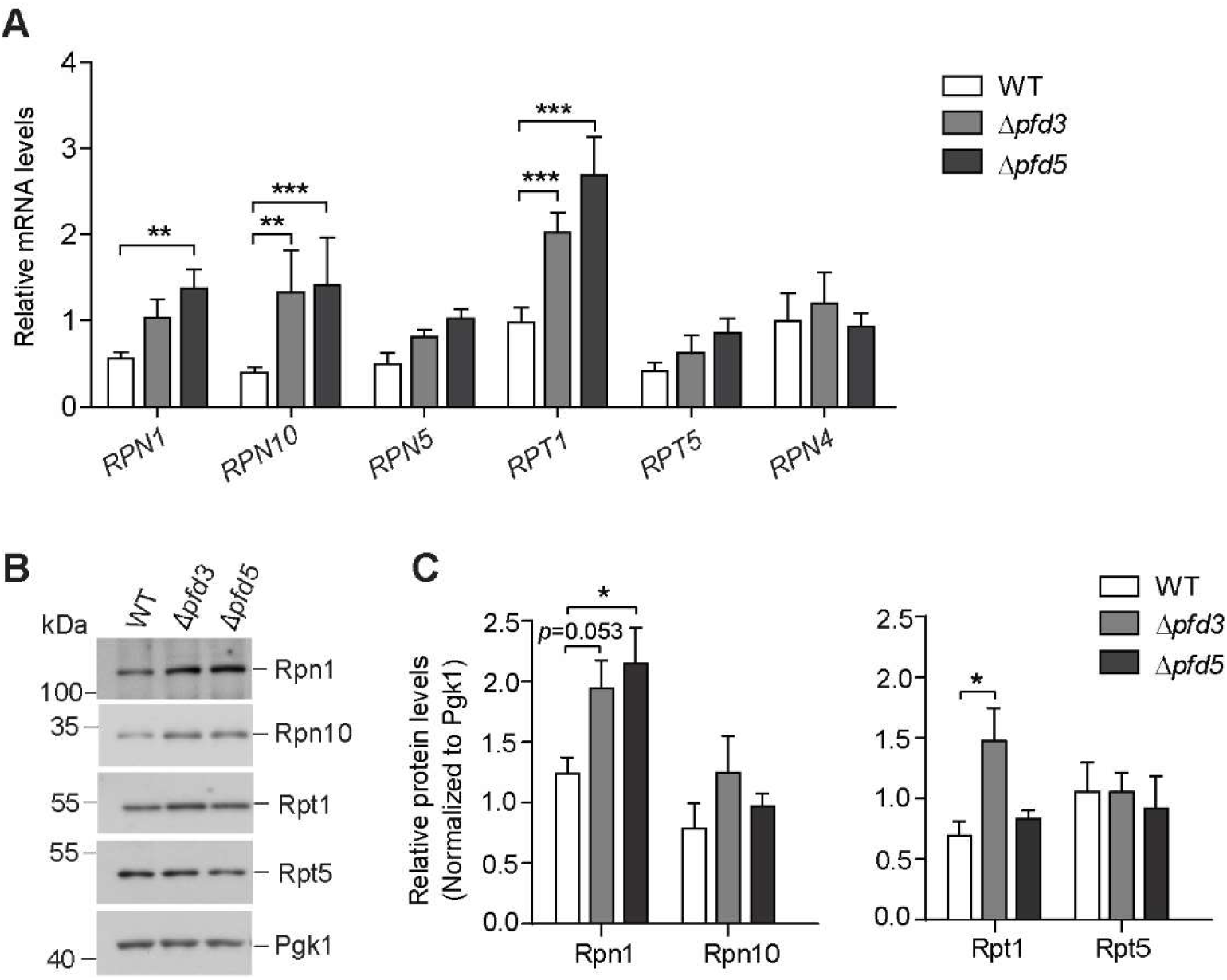
*PFD3* and *PFD5* deletions express higher levels of proteasome subunits. **A, B.** Wild type, Δ*pfd3* and Δ*pfd5* strains were grown on full growth medium containing glycerol at 25°C to logarithmic growth phase. **A.** The expression levels of indicated proteasome genes were assessed by qRT-PCR. The relative transcript levels of each gene to geometrical mean of three housekeeping genes including *ACT1, TDH1*, and *ALG9* were calculated. The data are represented as mean ± SEM, *n* ≥ 3. \*\*\**p* < 0.001, \*\**p* < 0.01. **B.** Total proteins were separated by SDS-PAGE and analysed by immunoblotting using antibodies against two non-ATPase subunits (Rpn1, Rpn10) and two ATPase subunits (Rpt1, Rpt5) of the regulatory particle. **C.** Quantification of western blot. Relative levels of proteasome subunits to Pgk1. The data are expressed as mean ± SEM, *n* ≥ 3. \**p* < 0.05. WT, wild type.

The expression of *RPN4*, the transcription factor responsible for the up-regulation of proteasome subunits, was unchanged in the deletions compared to wild-type cells, as expected, since it is regulated at the protein level by stabilisation due to insufficient proteasomal degradation (Ju et al., 2004; Xie & Varshavsky, 2001) (Figure 3A). We then examined the protein levels of the selected proteasome subunits. Protein levels were only partially consistent with transcript levels. Rpn1 levels were increased in prefoldin subunit deletions compared to wild type (Figure 3B, C). Rpn10, Rpt1 and Rpt5 subunits were not increased in the prefoldin subunit 5 deletion, but were also not decreased compared to the protein levels of wild-type cells. Taken together, our data suggest that Δ*pfd3* and Δ*pfd5* strains have a greater tendency to upregulate proteasome subunits and that deletion cells are not limited in the production of proteasome subunits.

### Prefoldin subunit deletions differ in accumulating ubiquitinated proteins

The increase in proteasome subunits at the transcript levels was likely triggered by insufficient proteosomal degradation capacity (Ju et al., 2004; Xie & Varshavsky, 2001). Consistent with previous studies, at physiological temperature we found that deletion of prefoldin subunits resulted in the accumulation of ubiquitinated protein species but not to the same extent. No accumulation was observed in Δ*pfd4* compared to wild-type cells. The accumulation of ubiquitinated proteins was highest in Δ*pfd3* and Δ*pfd5*. Cells lacking *PFD2* showed an intermediate phenotype (Fig. 4A, Supplementary Fig. 2A).

**Fig. 4.**
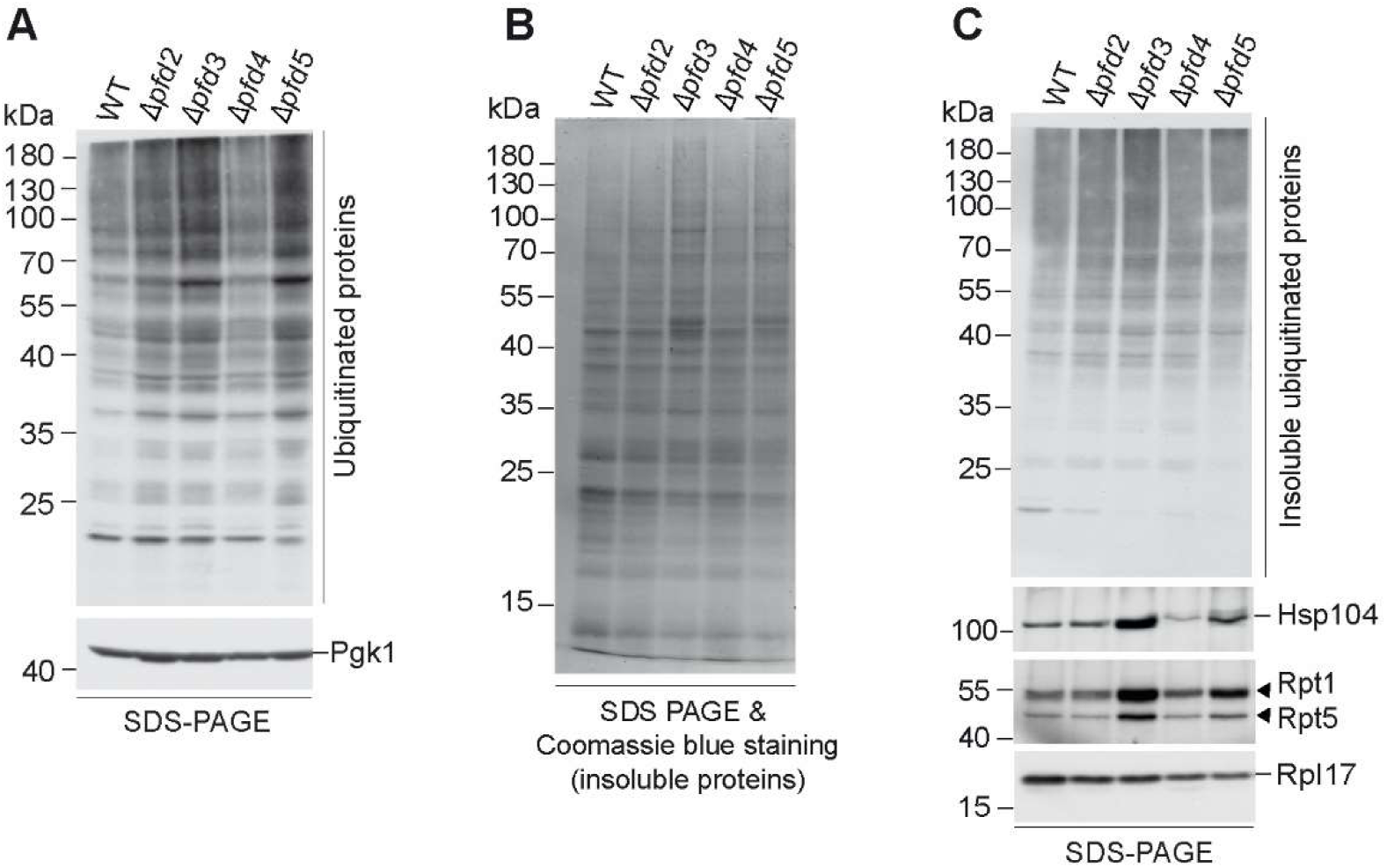
Degradation of ubiquitinated protein species is supported by prefoldin subunits. **A-C.** Wild type and indicated prefoldin single subunit deletion strains were cultured on full growth medium containing glycerol at 25°C. Protein extracts were prepared from equal OD_600_ units of cells. **A.** The same volume of protein extract was separated by SDS-PAGE and analysed by immunoblotting. **B.** Insoluble proteins were pelleted by ultracentrifugation and analysed by SDS-PAGE followed by coomassie blue staining. **C.** Insoluble proteins were separated by SDS-PAGE followed by immunoblotting using indicated antibodies. The experiments were performed in at least two biological repetitions. WT, wild type.

We tested whether prefoldin subunit deletions increase the accumulation of insoluble proteins at physiological temperature. Among the strains tested, only the Δ*pfd3* and Δ*pfd5* strains showed a shift towards higher molecular weight proteins in the insoluble protein fraction (Fig. 4B). The insoluble fractions were characterised by the presence of ubiquitinated protein species, but an increase compared to wild-type cells was only observed upon deletion of *PFD3* (Fig. 4C). Mass spectrometry analysis of the insoluble fractions of wild-type and Δ*pfd3* cells showed that similar classes of proteins accumulated in wild-type and deletion cells at similar levels (Supplementary Fig. 2B, C). Proteins in the insoluble fractions included ribosomal proteins, mitochondrial proteins and chaperones (Supplementary Fig. 2D). Thus, our data support a scenario in which ubiquitinated proteins are not efficiently degraded upon impairment of prefoldin function and such proteins may also accumulate in an insoluble protein fraction.

Targeted analysis by western blot showed that proteins with increased levels in the insoluble fraction of Δ*pfd3* and Δ*pfd5* cells compared to wild type included Hsp104, a protein disaggregase and marker for aggregates, but also the proteasome subunits Rpt1 and Rpt5. The Δ*pfd2* and Δ*pfd4* strains did not show accumulation of Hsp104, Rpt1 or Rpt5 compared to wild-type cells (Fig. 4C).

Thus, so far we showed that deletion of *PFD3* and *PFD5* resulted in the accumulation of ubiquitinated proteins, suggesting that both deletions caused proteasome defects to which cells adapt by increasing proteasome subunit levels. However, while these subunits are likely to be incorporated into the proteasome complex in Δ*pfd3*, cells lacking *PFD5* fail to fully assemble the proteasome. Interestingly, compared to wild type and other prefoldin subunit deletions, including Δ*pfd5*, we observed a greater tendency for a decrease in global translation in Δ*pfd3* cells (Supplementary Fig. 3A, B). A slowdown in protein synthesis has previously been shown to be beneficial for proteasome assembly (Nahar et al., 2019) and may potentially contribute to proteasome assembly in Δ*pfd3* cells.

### Heat stress-induced proteasome assembly is compromised upon loss of *PFD5*

Heat stress causes proteostatic stress, which increases the demand for degradation of misfolded proteins. We investigated the response of cells lacking *PFD3* or *PFD5* on proteasome assembly and activity under stress conditions. First, we measured proteasome chymotrypsin-like and caspase-like activities by an in-solution based approach using fluorogenic peptide substrates (Fig. 5A, B).

**Fig. 5.**
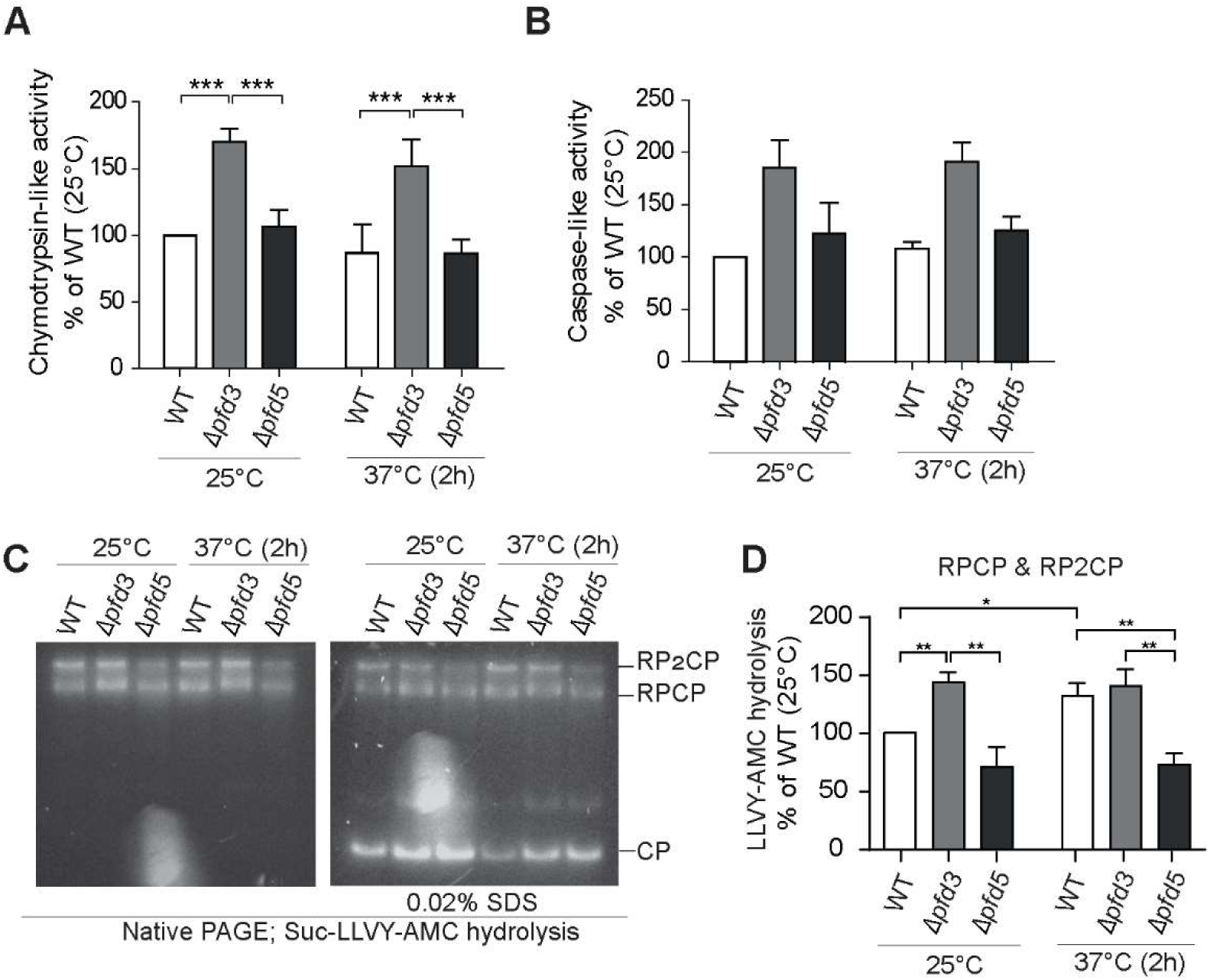
Heat-stress response is insufficient to activate proteasome activity in prefoldin 5 deletion. **A, B.** Wild type, Δ*pfd3* and Δ*pfd5* strains were grown on full growth medium containing glycerol at 25°C to logarithmic growth phase and then shifted to 37°C for 2 hours. Equal protein concentration were used for assessment of in-solution proteasome proteolytic activity using substrates specific for chymotrypsin-like, Suc-LLVY-AMC (A), and caspase-like, Ac-Nle-Pro-Nle-Asp-AMC (B), activities. The kinetics of substrate cleavage were assessed by measuring the fluorescence signal every 5 min during 1-2 hours by microplate reader. For each reaction, a linearized graph was plotted and the slop value, showing the kinetics of the reactions, was determined. The percentage of proteolytic activity for each strain to that in wild type (at 25 °C) was calculated and presented as bar charts. Data are expressed as mean ± SEM, *n* = 4 (A); *n* = 2 (B). \*\*\**p* < 0.001. **C.** Cells were grown on full growth medium containing glycerol at 25°C to logarithmic growth phase and then shifted to 37°C for 2 hours. Equal concentration of cell lysates were resolved by native-PAGE to assess the proteasome proteolytic activity by Suc-LLVY-AMC hydrolysis. **D.** The levels of RPCP and RP2CP shown in (C) were quantified and normalized to levels of those in wild type at 25 °C. Data are represented as mean ± SEM, *n* ≥ 3. \*\**p* < 0.01, \**p* < 0.05. WT, wild type.

The analysis revealed higher chymotrypsin-like and caspase-like activities in *PFD3* deletion cells compared to wild type, whereas total peptidase activity in Δ*pfd5* cells was similar to wild type. A similar pattern of proteolysis activity was observed under physiological and heat stress conditions. The proteasome in-solution assay does not distinguish between the peptidase activities of 26S or 20S proteasome. Thus, we applied the in-gel peptidase assay (Fig. 5C). It showed higher activity of RPCP and RP2CP complexes in Δ*pfd3* cells compared to wild-type cells, which is in agreement with the in-solution assay (Fig. 5C, D). As shown before, the peptidase activity of RPCP and RP2CP proteasome complexes was lower in the *PFD5* deletion strain compared with wild-type cells at 25°C (see also Fig 1A) but CP activity was increased. Heat stress increased the proteasome-peptidase activity of wild-type cells but not in Δ*pfd3* nor Δ*pfd5* cells (Fig. 5 C, D). We concluded that stress responses induced by high temperature could not compensate for the lack of prefoldin subunit function during proteasome assembly.

### Pfd5 deletion limits assembly of the Rpt-ring of the regulatory proteasome particle

We asked whether there was a limiting step in proteasome assembly in Δ*pfd5* cells. Using immunoblotting with the antibody against Rpt5, we observed several intermediate forms of assembly (Fig. 6A). Rpt5 is part of the base subcomplex, which contains six different AAA+ ATPases, Rpt1-6, which form a hetero-hexameric ring and form the molecular motor of the proteasome. The assembly chaperones Nas2, Nas6, Hsm3, Rpn14, and Adc17 have been shown to support assembly of the base subcomplex (Budenholzer et al., 2017). We investigated whether Pfd5 could act in a similar way to these assembly chaperones during base formation. Therefore, we made double deletions of *PFD5* with *HSM3* or *NAS6*. A growth assay showed a strong growth defect in the *PFD5;HSM3* double deletion under heat stress conditions (Supplementary Figure 4, 37°C). Analysis of in-gel peptidase activity showed a further decrease in activity of RP2CP particles in the double deletion of *PFD5;HSM3* (Figure 6B). The increased defect in activity was reflected in decreased levels of RP2CP (Fig. 6C).

**Fig. 6.**
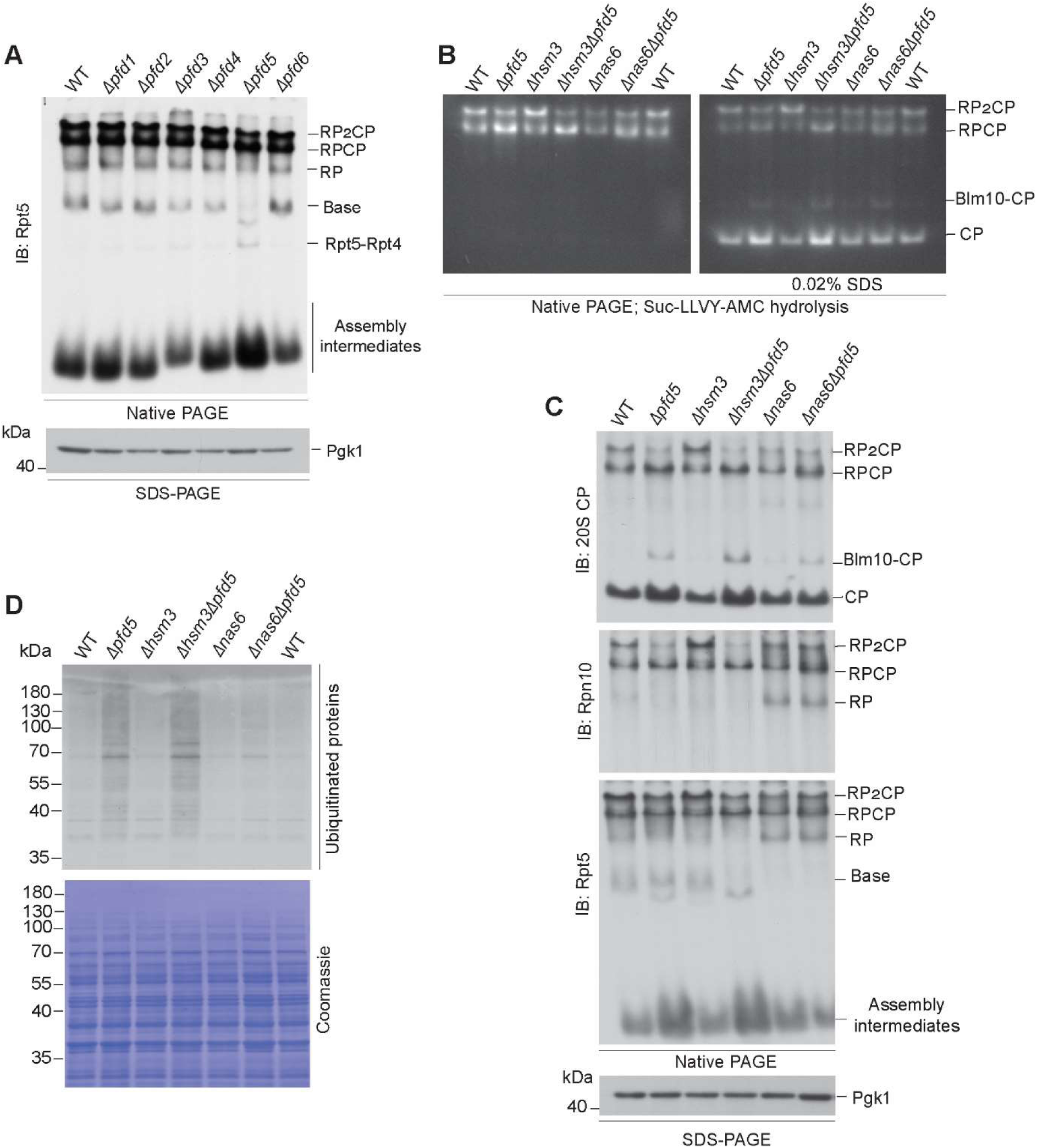
Pfd5 contributes to the assembly of the proteasomes Rpt-ring. **A.** Cells were grown on full growth medium containing glycerol at 25 °C to logarithmic growth phase. Equal concentration of cell lysates were resolved by native-PAGE and SDS-PAGE and analysed by western blot. **B, C.** Equal concentrations of cell lysates were resolved by native-PAGE to assess the proteasome proteolytic activity by Suc-LLVY-AMC hydrolysis (B) and by immunoblotting (C). **D.** Total cell lysates were separated on SDS-PAGE and analysed by western blot using anti-ubiquitin antibody. Coomassie staining is presented as loading control.

In contrast, deletion of *PFD5* in *nas6*Δ reduced the growth of the *nas6*Δ strain under heat stress conditions, but not beyond the growth of *pfd5*Δ (Supplementary Figure 4). Surprisingly, deletion of *NAS6* in *pfd5*Δ improved proteasome assembly, reduced assembly intermediates detected by anti-Rpt5 antibody and decreased CP levels (Fig. 6C).The increased levels of assembled proteasomes were reflected in proteasome activity, which was similar to wild-type cells (Fig. 6B). Finally, we observed the consequences of the altered proteasome assembly in the double deletions. Consistent with our observations, the accumulation of polyubiquitinated protein species was further increased in the *PFD5;HSM3* double deletion, but decreased in the *PFD5;NAS6* double deletion compared with *pfd5*Δ only (Fig. 6D). Thus, our data suggest that Pfd5 is required for certain steps of base subcomplex assembly, possibly in parallel with the function of Hsm3.

## Discussion

This work consolidates the role of the co-chaperone prefoldin within the proteostasis network by demonstrating that the absence of a functional prefoldin complex impairs the stability and/or limits synthesis of newly produced 26S proteasome, resulting in the accumulation of ubiquitinated protein species. In particular, our work has shown that deletion of the prefoldin subunits Pfd3 and Pfd5 increases the levels of ubiquitinated proteins under physiological growth conditions suggesting inefficient degradation of ubiquitinated substrates. Our data are consistent with previous observations showing that si-RNA-mediated knockdown of *PFDN5* increased ubiquitinated proteins in HeLa cells following chemical inhibition of the proteasome (Abe et al., 2013). In addition, polyubiquitinated proteins accumulated in cell lines derived from a mouse model with a missense mutation in *Pfdn5* (Abe et al., 2013). Furthermore, the prefoldin complex co-localised with insoluble aggregates in cell culture, suggesting that the co-chaperone performs protein quality control to reduce protein aggregates, including those implicated in neurodegenerative diseases. (Abe et al., 2013; Sorgjerd et al., 2013; Tashiro et al., 2013). While these observations are noteworthy, the mechanism by which prefoldin regulates the levels of ubiquitinated proteins and insoluble aggregates was unknown. Here we propose an active role for the prefoldin complex in protein quality control by contributing to the formation of the proteasome complex, rather than simply impairing protein folding.

Prior to degradation by the proteasome complex, the RP identifies the polyubiquitinated targets in the cell through its intrinsic ubiquitin receptors, Rpn1, Rpn10 and Rpn13 (Saeki, 2017; Verma et al., 2004). Simultaneous recognition of the unstructured initiation region of substrates by the Rpt ring ensures substrate engagement (Jonsson et al., 2022; Prakash et al., 2004). Subsequently, the engaged substrate undergoes deubiquitination by Rpn11 (Verma et al., 2002), mechanical unfolding and translocation into the proteolytic chamber by the Rpt ring (Bard et al., 2019). Structural defects in the RP prevent substrate processing and final degradation of polyubiquitinated substrates by the proteasome (Matyskiela et al., 2013; Wehmer et al., 2017). It has been shown that, in contrast to CP, RP modules have fragile structures and their assembly requires the proteasome assembly chaperones Nas6, Rpn14, Hsm3, Nas2 and Adc17 (Hanssum et al., 2014; Kaneko et al., 2009; Le Tallec et al., 2009; Park et al., 2009; Roelofs et al., 2009; Saeki et al., 2009; Thompson et al., 2009). Inefficient RP assembly due to lack of assembly checkpoints leads to the accumulation of ubiquitinated protein species that are not processed by RP and ultimately remain undegraded. The presence of proteasome complex assembly intermediates detected by using antibodies against RP proteins (Rpn10, Rpt5) in Δ*pfd5* cells suggests that Pfd5 is important for the assembly and integrity of proteasome complexes. Prefoldin acts downstream of the translation machinery. A plausible explanation could be that prefoldin, and Pfd5 in particular, is involved in the folding of newly folded polypeptides of proteasome subunits. Certainly, protein folding requires an ATP-driven chaperone such as the type II chaperonin TRiC/CCT. Indeed, Cct subunits have been identified as proteasome-interacting proteins in a mass spectrometry approach and the authors speculate that the Cct complex functions in assisting protein degradation through its interaction with the proteasome rather than being a substrate itself (Guerrero et al., 2008). On the other hand, the first step in the formation of the Rpt ring requires the formation of specific pairs of ATPases, namely Rpt1-Rpt2, Rpt3-Rpt6 and Rpt4-Rpt5. It has been shown that the nascent polypeptides of Rpt1 and Rpt2 assemble co-translationally and this was supported by the Hsm3 chaperone, but its presence was not essential for dimer formation (Panasenko et al., 2019). This suggests either the involvement of other chaperones or spontaneous formation of the heterodimer. The increased defect in proteasome assembly in the *PFD5;HSM3* double deletion suggests that Pfd5 may be important for Rpt dimer formation. However, to our knowledge, structural analysis did not detect Pfd5 nor prefoldin complex in complex with proteasome intermediates and therefore the interaction could be considered transient. A similar observation has previously been made with Adc17, which binds transiently to Rpt6 and has been proposed to assist in the formation of Rpt3 and Rpt6 dimer, but dissociates prior binding of Nas6 to Rpt3 forming the Nas6-Rpt3-Rpt6-Rpn14 module. Furthermore, single deletions of the Rpt ring chaperones are viable but double deletions can show compromised growth upon proteotoxic stress. This is in agreement with our observation where the double deletion of *PFD5* and *HSM3* showed strong growth defect at heat stress conditions.

The hierarchy of assembly events from ATPase dimers to the Rpt ring structure is not fully understood. The rescue of the Pfd5-dependent proteasome defect by deletion of *NAS6* is somewhat surprising, but high-throughput analysis previously revealed positive genetic interactions between *pfd5Δ* and deletion of *RPT4*, *RPN10* and *RPN12* (Costanzo et al., 2016; Wilmes et al., 2008). However, the finding of epistatic genetic interactions suggests a sequential requirement for Pfd5 during the assembly process. Further epistasis analysis will be necessary to reveal the functional relationships between the Pfd5 and Rpt ring chaperones.

In conclusion, by discovering a non-canonical function of Pfd5 in proteasome assembly, our work adds to the repertoire of factors that are canonically important in controlling the homeostatic levels of proteasomes.

## Materials and methods

### Yeast strains and growth conditions

*Saccharomyces cerevisiae* strains used in this study were derivatives of BY4741 (MATa, his3Δ1; leu2Δ0; met15Δ0; ura3Δ0) purchased from Euroscarf (http://www.euroscarf.de). The deletion of *PFD* gene in *pfd*Δ strains, *hsm3*Δ and *nas6*Δ were confirmed by amplification of kanMx cassette and sequencing of deletion specific barcode regions. Yeast strain carrying the double deletion of *HSM3* or *NAS6* with *PFD5* were generated by the replacement of *HSM3* or *NAS6* genes with *HIS3* cassette amplified from pRS303 vector in a BY4741 strain carrying a single *PFD5* deletion. Deletion of *HSM3*, *NAS6* and *PFD5* was checked by PCR and sequencing. Yeast strains used and generated in this study are listed in Table 1.

**Table 1.** *Saccharomyces cerevisiae* BY4741 strains used in this study.

| Strain name | Genotype<sup>1</sup> | Source |
| --- | --- | --- |
| *Δpfd1* | *Δpfd1::KanMX4* | Euroscarf |
| *Δpfd2* | *Δpfd2::KanMX4* | Euroscarf |
| *Δpfd3* | *Δpfd3::KanMX4* | Euroscarf |
| *Δpfd4* | *Δpfd4::KanMX4* | Euroscarf |
| *Δpfd5* | *Δpfd5::KanMX4* | Euroscarf |
| *Δpfd6* | *Δpfd6::KanMX4* | Euroscarf |
| *Δhsm3* | *Δhsm3::KanMX4* | Horizon Discovery/<br>Yeast knockout collection |
| *Δnas6* | *Δnas6::KanMX4* | Horizon Discovery/<br>Yeast knockout collection |
| *Δpfd5Δhsm3* | *Δpfd5::KanMX4 Δhsm3::HIS3* | This study |
| *Δpfd5Δnas6* | *Δpfd5::KanMX4 Δnas6::HIS3* | This study |

<sup>1</sup>All BY4741 strains are haploid with the background *MATa his3Δ1 leu2Δ0 met15Δ0 ura3Δ0*.

The yeast strains were grown on full medium, YP medium (1% yeast extract and 2% bacto peptone), or minimal medium (0.67% yeast nitrogen base without amino acids and 0.079% CSM amino acid mix) supplemented with 2% glucose or 3% glycerol as fermentable and respiratory carbon sources, respectively. The strains were generally grown at 25°C and when indicated shifted to restrictive temperature (37°C) for 2 hours.

### DNA manipulation and generation of constructs

To rescue *PFD5* expression in Δ*pfd5* strains, *PFD5* open reading frame along with 398 bp promoter region and 202 bp of the downstream sequence was amplified from genomic DNA of wild-type strain (BY4741) using specific primers with at least 15 bp homology extension to the linearized vector ends. The amplified fragments were verified and subcloned into the pRS415 vector by the ligation-independent cloning method as described previously (Jeong et al., 2012). The sequence of primers used for cloning is listed in Table 2. All generated constructs were verified by DNA sequencing before the transformation. The resulting transformants were grown on minimal selective medium supplemented with 2% glucose or 3% glycerol as required.

**Table 2.**
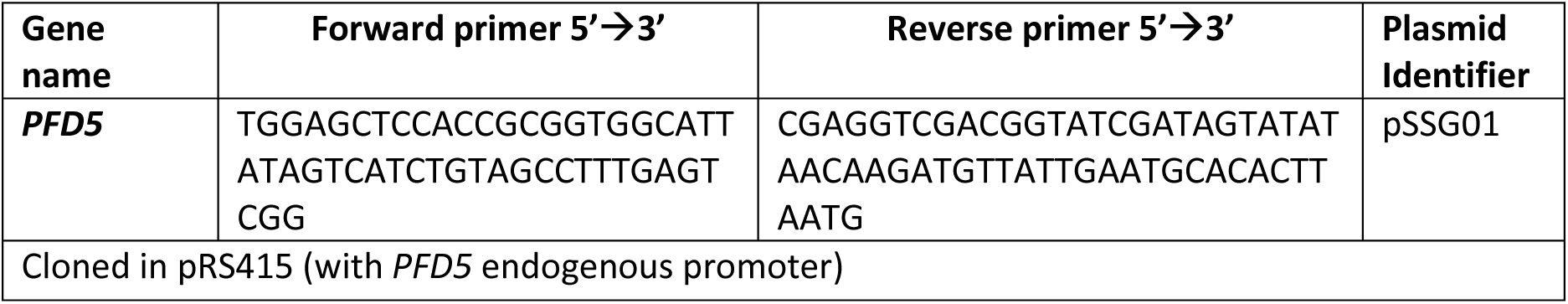
The construct generated in this study.

### Total protein isolation and immunoblot analysis

Total yeast protein was isolated from 2.5 OD_600_ units of exponentially grown yeast cells (0.6-0.8 OD_600_) according to a previously described protocol (Yaffe & Schatz, 1984). Proteins were solubilized in Laemmli buffer containing 50 mM DTT. Samples were denatured at 65°C for 15 min and a sample volume corresponding to 0.1-0.2 OD_600_ unit was resolved by SDS-polyacrylamide gel electrophoresis (SDS-PAGE). Western blot technique was carried out using standard protocols. Custom primary antibodies were generated in rabbits to detect yeast Rpl17, and Rpn1. Following commercially available antibodies were used: anti-Pgk1 (459250; Thermo Fisher Scientific), anti-Hsp104 (ADI-SPA-1040-D; Enzo Life Sciences), anti-Ubiquitin-P4D1 (sc-8017; Santa cruz Biotechnology), anti-Rpt1 (BML-PW8255; Enzo Life Sciences), anti-Rpt5 (BML-PW8245, Enzo Life Sciences), anti-Rpn5 (ab79773; Abcam), anti-Rpn10 (ab98843; Abcam), anti-proteasome 20S core subunits (BML-PW9355; Enzo Life Sciences), anti-DNP antibody (MAB2223, Merck), anti-rabbit IgG secondary antibody (A9169; Sigma-Aldrich), anti-Mouse IgG secondary antibody (A4416-1ML; Sigma-Aldrich).

### RNA Isolation and quantitative RT-PCR

Total RNA was isolated from 20 OD_600_ units of logarithmic grown yeast cells using hot acid-phenol and SDS method as described previously (Schmitt et al., 1990). About 40 ng of total RNA was used to synthesize cDNA by using QuantiTect reverse transcriptase kit (205313; Qiagen) according to the manufacturer protocol. The quantitative reverse transcription PCR (qRT-PCR) was performed using RT PCR Mix SYBR C (2008-100C; A&A Biotechnology) and specific primers (Table 3) on the Roche LightCycler 480 System (Basel, Switzerland). The reaction program includes initial denaturation for 5 min at 95 °C, followed by 40 cycles of 30 s (95 °C), 20 s (58 °C), and 20 s (72°C). A standard curve was generated by analysing serial dilutions of cDNA for each pairs of primers. In addition to samples and standards, template-negative and RT-negative controls were included to the measurements in parallel. The relative expression of a given gene to the geometric mean of three reference genes including *ACT1*, *ALG9*, and *TDH1* was calculated and represented on a column graph.

**Table 3.**
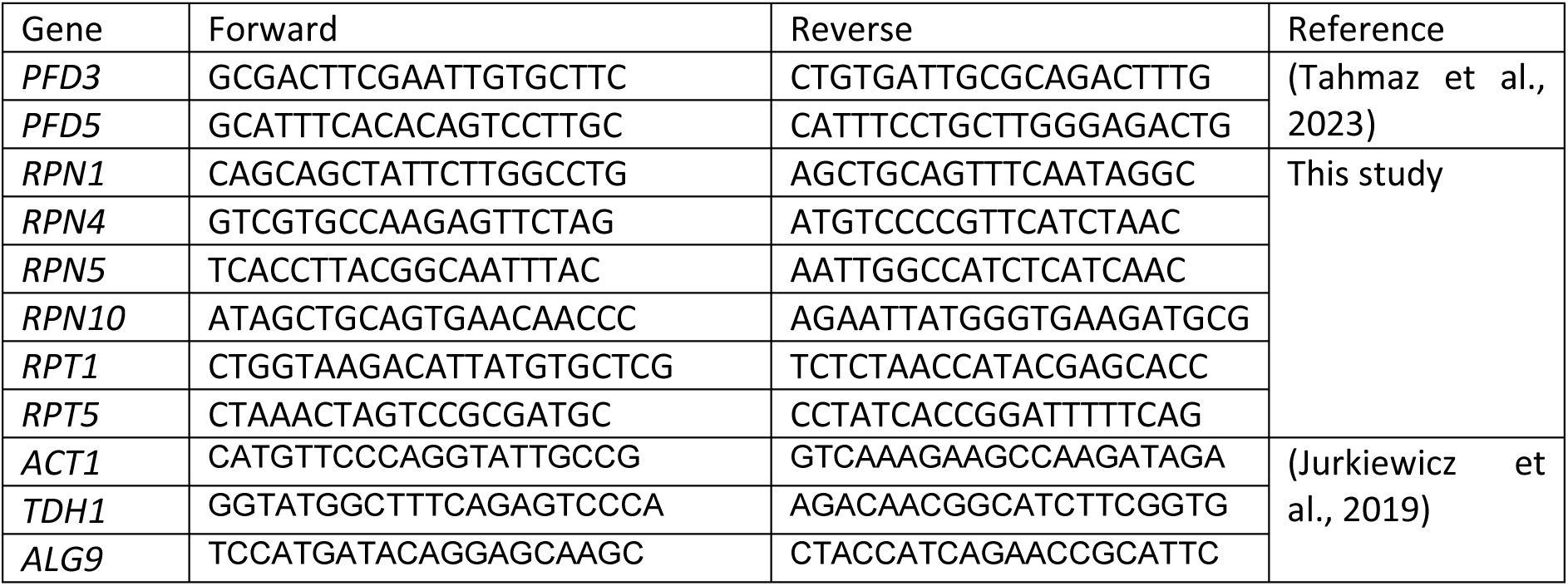
Sequence of primers (5’-3’) used for qRT-PCR.

### Protein aggregation assay

About 10 OD_600_ units of yeast cells at logarithmic growth phase (OD_600_ = 0.6-0.8) were pelleted, washed with water and resuspended in 400 μl of lysis buffer (30 mM Tris-HCl pH 7.4, 150 mM NaCl, 20 mM KCl, 5 mM EDTA pH 8.0 and 0.5 mM PMSF). Cells were broken by addition of 200 μl of glass beads and shaking on Cell Disruptor Genie™ (Thermo Fisher Scientific) for 10 min at 4°C. After the beads settled down, 1% [v/v] Triton X-100 was added to the lysate followed by incubation for 10 min at 4°C. Cell extract was clarified by centrifugation (4000xg, 10 min, 4°C), and subsequently fractionated into supernatant and pellet of aggregated proteins by ultracentrifugation (125000xg, for 1 hour at 4°C). The pellet was resuspended in 65 µl of urea sample buffer (6M urea, 6% SDS, 125mM Tris-HCl pH 6.8, and 0.01% Bromophenol Blue). Proteins from total and supernatant fractions were isolated following TCA precipitation (final concentration of 10%), and resuspended in 65 µl of the urea sample buffer.

For preparation of aggregated proteins for mass spectrometry analysis, 10 mM Chloroacetamide was added to the lysis buffer. After ultracentrifugation, pelleted proteins were washed with 200 µl of methanol to remove Triton X-100. The pellet was washed with acetone, and resuspended in 50 µl of solubilization buffer (8 M Urea and 100 mM TRIS pH 8.5) containing 0.1% of ProteaseMAX^TM^ surfactant (V2072; promega, in 50mM ammonium bicarbonate) and 10 mM TCEP. The suspension was sonicated on ice (3 times, each time 3 sec. with 10 sec. break after each sonication) and subjected to further processing before MS analysis (see section “Preparation of samples for proteomics analysis”).

### Preparation of samples for proteomics analysis

MS analysis of insoluble proteins was performed using the FASP method described previously (Wisniewski et al., 2009) with few modifications. Final washes and protease digestion were performed in 50 mM ammonium bicarbonate containing 0.1% of ProteaseMAX^TM^ surfactant (V2072; Promega). Digestion was carried out overnight using trypsin (Promega) 1:100 protease/protein ration at 37 °C. Peptides were eluted from spin filters twice by additions of 100ul of 50 mM ammonium bicarbonate. Eluted peptide concentration was measured by Direct Detect^®^ spectrometer (Merck). Samples were dried by speedvac and resuspended in appropriate volume of 0.1% formic acid (FA) in water.

### Mass spectrometry and data analysis

Chromatographic separation was performed on an Easy-Spray Acclaim PepMap column 50 cm length × 75 µm inner diameter (Thermo Fisher Scientific) at 55°C by applying 90 min acetonitrile gradients in 0.1% aqueous formic acid at a flow rate of 300 nl/min. An UltiMate 3000 nano-LC system was coupled to a Q Exactive HF-X mass spectrometer via an Easy-Spray source (all Thermo Fisher Scientific). The Q Exactive HF-X was operated in data-dependent mode with survey scans acquired at a resolution of 120,000 at m/z 200. Up to 12 of the most abundant isotope patterns with charges 2-5 from the survey scan were selected with an isolation window of 1.3 m/z and fragmented by higher-energy collision dissociation (HCD) with normalized collision energies of 27, while the dynamic exclusion was set to 30 s. The maximum ion injection times for the survey scan and the MS/MS scans (acquired with a resolution of 15,000 at m/z 200) were 45 and 96 ms, respectively. The ion target value for MS was set to 3 × 1e6 and for MS/MS to 1e5, and the intensity threshold for MS/MS was set to 1e4.

Proteins were identified and quantified with LFQ algorithm using MaxQuant (version 2.3.0.0) and UNIPROT *Saccharomyces cerevisiae* reference proteome UP000002311 as search database. Following MaxQuant parameters were applied; Digestion: Trypsin/P, maximum 2 missed cleavages, Fixed modifications – Carbamidomethyl (C), Variable modifications – Oxidation (M), Acetyl (Protein N-ter), FDR 0.01. All other parameters were set to default. Further analysis was performed with Perseus software (version 2.0.9.0.). Data were cleaned and only peptides with Razor + unique peptides > 1 were maintained. Data were log2 transformed and missing values were imputed from normal distribution per column (width 0.3, down shift 1.8). Median of log2 transformed LFQ values of four biological replicates per condition was calculated and displayed as heat map.

### Proteasome in-solution activity assay

About 20 OD_600_ units of exponentially grown yeast cells (0.6-0.8 OD_600_) were harvested and washed with water. The cell pellet was resuspended in 600 µl of ice-cold lysis buffer (50 mM Tris-HCl pH 7.4, 10 mM MgCl2, 10% glycerol) containing freshly added 2 mM ATP, 2 mM PMSF, and 1 mM DTT. About 400 µl glass beads were added and cells were broken at 4°C by Cell Disruptor Genie™ (Thermo Fisher Scientific) for 7 min. The lysate was cleared by centrifugation (20.000xg, 15 min, 4°C) and protein concentration was determined by Direct Detect^®^ spectrometer (Merck). The proteasome activity was measured using a 96-well plate. Each well contained 50 µg of protein extract and 100 µM of fluorogenic peptide substrate in a total volume of 200 µl. Peptide substrates Suc-LLVY-AMC (I-1393; Bachem) and Ac-Nle-Pro-Nle-Asp-AMC (I-1850; Bachem,) were used to assess proteasome chymotrypsin-like activity and caspase-like activity, respectively. Negative control samples were prepared by mixing 50 µg of protein extract and DMSO (equal volume as peptide substrate). To test the specificity of proteasome cleavage, 50 µg of protein extract was treated with 50 µM MG132, a potent proteasome inhibitor, before adding peptide substrate. Each test and control samples were prepared at least in duplicates. The reaction kinetics was assessed by measuring fluorescence signal (380 nm excitation wavelength, 460 nm emission wavelength) every 5 minutes for a duration of 1-2 hours, by SpectraMax^®^ iD3 Multi-Mode Microplate Reader (Molecular Devices). Proteasome activity was calculated based on the line graph slope value, after the deduction of background and non-proteasome cleavage of a substrate (controls) from the tested samples.

### In-gel overlay proteasome assay

About 30 OD_600_ units of exponentially grown yeast cells (0.6-0.8 OD_600_) were harvested and washed with water. The cell pellet was resuspended in 500 µl of cold lysis buffer containing 50 mM Tris-HCl pH 7.4, 10 mM MgCl2, 250 mM sucrose, 2 mM ATP, 2 mM PMSF and 1 mM DTT. The cell lysate was prepared and protein concentration was measured by Direct Detect^®^ spectrometer (Merck). An equal amount of proteins (50 µg) were separated on 4.5% polyacrylamide native gel containing 2.5% sucrose, 1 mM ATP, 1M DTT, 10% APS, (1:1000) TEMED, and 1X native running buffer (90 mM Tris base, 90 mM boric acid, 5 mM MgCl2, 0.5mM EDTA and 1 mM fresh ATP) for 1h (20 mA), 3 h (28 mA) and 1h (22 mA), in the cold room. As a loading control, part of the lysate was subjected to SDS-PAGE and Pgk1 levels were analyzed. After completing the electrophoresis, the native gel was either subjected to analysis of proteasome composition by immunoblotting or used for in-gel proteasome activity assay as described previously (Elsasser et al., 2005; Yazgili et al., 2021). For the latter purpose, the gel was incubated in freshly prepared reaction buffer (50 mM Tris-HCl pH 7.4, 10 mM MgCl2, 250 mM sucrose, 1mM ATP) containing 50 µM peptide substrate Suc-LLVY-AMC and incubated at 37°C for 30 minutes in dark. The gel was imaged by exposing it to UV light (380 nm excitation, 460 nm emission) in the Alliance Q9 Imager (UVITEC Cambridge). The activity of the 20S core complex was stimulated by adding 0.02% SDS to the reaction buffer.

### Carbonylation assay

Crude cell extracts were prepared by incubation of 18 OD_600_ units of yeast cells in lysis buffer (20 mM Tris-HCl pH 7.4, 200 mM NaCl, 0.5% Triton X-100, 1 mM PMSF). Cells were disrupted with glass beads. The supernatant was cleared by centrifugation at 3,000 × g for 1 min. To precipitate DNA in the protein lysate, streptomycin diluted in 50 mM 4-(2-hydroxyethyl)-1-piperazine ethanesulfonic acid (HEPES) pH 7.0 was added at a final concentration of 1% and incubated for 15 min on ice followed by centrifugation at 6,000 × g for 10 min at 4°C. Four volumes of 10 mM 2,4-dinitrophenylhydrazine (DNPH) diluted in 2 N hydrochloric acid (HCl) was added to the lysate. The samples were incubated for 1 h at RT protected from light. Proteins were precipitated by adding TCA to a final concentration of 10%, followed by incubation on ice for 30 min and centrifugation at 20,000 × g for 15 min at 4°C. The pellets were washed three times with an ice-cold 1:1 (v/v) mixture of ethanol and ethyl acetate and resuspended in 8M Urea. Carbonylated proteins were detected by immunoblotting using anti-DNP antibody.

### ROS measurement

ROS levels were assessed according to the previously described method (Topf et al., 2018) with minor modifications. Two OD_600_ units of yeast cells were collected and washed with PBS (137 mM NaCl, 12 mM phosphate, 2.7 mM KCl, pH 7.4). Yeast cells were resuspended in PBS containing 5µM dihydroethidium (Sigma cat no. D7008). The following controls were prepared for each condition; PBS only, yeast cells resuspended in PBS without dye, and PBS only with dye. Samples were incubated for 40 minutes at room temperature in the dark prior to measurement. Fluorescence signal was measured at an excitation wavelength of 535 nm and an emission wavelength of 635 nm using a plate fluorometer (SpectraMax iD3, Molecular Devices). Fluorescence signals were collected at 5-minute intervals for 2 hours. Fluorescence units from each sample were corrected for autofluorescence and autoxidation of the dye in PBS. Corrected data are presented as mean ± SEM of 5 biological replicates.

### Translation assay

To assess the levels of newly synthesised proteins, yeast cells were grown on minimal medium containing glycerol. One OD_600_ unite of exponentially grown cells were washed and incubated in minimal medium lacking Met amino acid, for 20 minutes. Subsequently, proteins were radiolabeled using [^35^S]-labelled methionine (SRM-01H, Hartmann Analytic) at a final concentration of 6 µCi/ml for 1h. Yeast cells were washed with ddH_2_O and subjected to protein isolation followed by SDS-PAGE. Radioactive signals were captured by phosphor screens, which was scanned using Fujifilm FLA-7000 (GE Healthcare). Images of three independent experiments were quantified and data displayed on a column graph.

### Statistical analysis

To determine statistical difference of data obtained from RT-qPCR and in-solution peptidase activity one-way analysis of variance (ANOVA) followed by Tukey’s post hoc test was performed. For quantification of densitometry after western blot and fluorescent signal after in-gel peptidase activity the unpaired, two-sided Student’s *t*-test was applied.

## Acknowledgments

We thank Monika Stasiak and Katarzyna Jonak for help with the generation of double deletion strains and Magdalena Boguta for comments on the manuscript. Proteomics analysis of insoluble proteins was performed in the proteomics core facility of the International Institute of Molecular Mechanisms and Machines Polish Academy of Sciences (IMol PAS). SSG and UT acknowledge support by COST action ProteoCure CA20113. This study was funded by National Science Centre Poland (grant no. 2018/31/B/NZ1/02401). The funders had no role in the study design, data collection and analysis, or preparation of the manuscript.

## Author contributions

SSG performed experiments and analyzed the data. KD prepared samples for mass spectrometry and analysed data. UT conceptualized the study, analyzed the data, and provided supervision and funding for the study. SSG and UT wrote the manuscript. All authors approved the final version of the manuscript.

## Data availability

The materials that were generated within this work are available upon request from the corresponding author (U.T.).

## Competing interests

The authors declare that they have no competing interests.

## Notes

### Competing Interest Statement

The authors have declared no competing interest.

## References

Abe, A., Takahashi-Niki, K., Takekoshi, Y., et al. (2013). Prefoldin plays a role as a clearance factor in preventing proteasome inhibitor-induced protein aggregation. J Biol Chem, 288(39), 27764–27776. 10.1074/jbc.M113.476358

Bard, J. A. M., Bashore, C., Dong, K. C., et al. (2019). The 26S Proteasome Utilizes a Kinetic Gateway to Prioritize Substrate Degradation. Cell, 177(2), 286–298 e215. 10.1016/j.cell.2019.02.031

Bard, J. A. M., Goodall, E. A., Greene, E. R., et al. (2018). Structure and Function of the 26S Proteasome. Annu Rev Biochem, 87, 697–724. 10.1146/annurev-biochem-062917-011931

Budenholzer, L., Cheng, C. L., Li, Y., et al. (2017). Proteasome Structure and Assembly. J Mol Biol, 429(22), 3500–3524. 10.1016/j.jmb.2017.05.027

Costanzo, M., VanderSluis, B., Koch, E. N., et al. (2016). A global genetic interaction network maps a wiring diagram of cellular function. Science, 353(6306). 10.1126/science.aaf1420

Dange, T., Smith, D., Noy, T., et al. (2011). Blm10 protein promotes proteasomal substrate turnover by an active gating mechanism. J Biol Chem, 286(50), 42830–42839. 10.1074/jbc.M111.300178

Dong, Y., Zhang, S., Wu, Z., et al. (2019). Cryo-EM structures and dynamics of substrate-engaged human 26S proteasome. Nature, 565(7737), 49–55. 10.1038/s41586-018-0736-4

Elsasser, S., Schmidt, M., & Finley, D. (2005). Characterization of the proteasome using native gel electrophoresis. Methods Enzymol, 398, 353–363. 10.1016/S0076-6879(05)98029-4

Geissler, S., Siegers, K., & Schiebel, E. (1998). A novel protein complex promoting formation of functional alpha- and gamma-tubulin. EMBO J, 17(4), 952–966. 10.1093/emboj/17.4.952

Gestaut, D., Roh, S. H., Ma, B., et al. (2019). The Chaperonin TRiC/CCT Associates with Prefoldin through a Conserved Electrostatic Interface Essential for Cellular Proteostasis. Cell, 177(3), 751–765 e715. 10.1016/j.cell.2019.03.012

Glickman, M. H., Rubin, D. M., Coux, O., et al. (1998). A subcomplex of the proteasome regulatory particle required for ubiquitin-conjugate degradation and related to the COP9-signalosome and eIF3. Cell, 94(5), 615–623. 10.1016/s0092-8674(00)81603-7

Guerrero, C., Milenkovic, T., Przulj, N., et al. (2008). Characterization of the proteasome interaction network using a QTAX-based tag-team strategy and protein interaction network analysis. Proc Natl Acad Sci U S A, 105(36), 13333–13338. 10.1073/pnas.0801870105

Hanssum, A., Zhong, Z., Rousseau, A., et al. (2014). An inducible chaperone adapts proteasome assembly to stress. Mol Cell, 55(4), 566–577. 10.1016/j.molcel.2014.06.017

Herranz-Montoya, I., Park, S., & Djouder, N. (2021). A comprehensive analysis of prefoldins and their implication in cancer. iScience, 24(11), 103273. 10.1016/j.isci.2021.103273

Jeong, J. Y., Yim, H. S., Ryu, J. Y., et al. (2012). One-step sequence- and ligation-independent cloning as a rapid and versatile cloning method for functional genomics studies. Appl Environ Microbiol, 78(15), 5440–5443. 10.1128/AEM.00844-12

Jonsson, E., Htet, Z. M., Bard, J. A. M., et al. (2022). Ubiquitin modulates 26S proteasome conformational dynamics and promotes substrate degradation. Sci Adv, 8(51), eadd9520. 10.1126/sciadv.add9520

Ju, D., Wang, L., Mao, X., et al. (2004). Homeostatic regulation of the proteasome via an Rpn4-dependent feedback circuit. Biochem Biophys Res Commun, 321(1), 51–57. 10.1016/j.bbrc.2004.06.105

Jurkiewicz, A., Lesniewska, E., Ciesla, M., et al. (2019). Inhibition of tRNA Gene Transcription by the Immunosuppressant Mycophenolic Acid. Mol Cell Biol, 40(1). 10.1128/MCB.00294-19

Kaneko, T., Hamazaki, J., Iemura, S., et al. (2009). Assembly pathway of the Mammalian proteasome base subcomplex is mediated by multiple specific chaperones. Cell, 137(5), 914–925. 10.1016/j.cell.2009.05.008

Le Tallec, B., Barrault, M. B., Guerois, R., et al. (2009). Hsm3/S5b participates in the assembly pathway of the 19S regulatory particle of the proteasome. Mol Cell, 33(3), 389–399. 10.1016/j.molcel.2009.01.010

Leroux, M. R., Fandrich, M., Klunker, D., et al. (1999). MtGimC, a novel archaeal chaperone related to the eukaryotic chaperonin cofactor GimC/prefoldin. EMBO J, 18(23), 6730–6743. 10.1093/emboj/18.23.6730

Liang, J., Xia, L., Oyang, L., et al. (2020). The functions and mechanisms of prefoldin complex and prefoldin-subunits. Cell Biosci, 10, 87. 10.1186/s13578-020-00446-8

Martin-Benito, J., Boskovic, J., Gomez-Puertas, P., et al. (2002). Structure of eukaryotic prefoldin and of its complexes with unfolded actin and the cytosolic chaperonin CCT. EMBO J, 21(23), 6377–6386. 10.1093/emboj/cdf640

Mathieu, C., Pappu, R. V., & Taylor, J. P. (2020). Beyond aggregation: Pathological phase transitions in neurodegenerative disease. Science, 370(6512), 56–60. 10.1126/science.abb8032

Matyskiela, M. E., Lander, G. C., & Martin, A. (2013). Conformational switching of the 26S proteasome enables substrate degradation. Nat Struct Mol Biol, 20(7), 781–788. 10.1038/nsmb.2616

Meyer-Schwesinger, C. (2019). The ubiquitin-proteasome system in kidney physiology and disease. Nat Rev Nephrol, 15(7), 393–411. 10.1038/s41581-019-0148-1

Nahar, A., Fu, X., Polovin, G., et al. (2019). Two alternative mechanisms regulate the onset of chaperone-mediated assembly of the proteasomal ATPases. J Biol Chem, 294(16), 6562–6577. 10.1074/jbc.RA118.006298

Panasenko, O. O., Somasekharan, S. P., Villanyi, Z., et al. (2019). Co-translational assembly of proteasome subunits in NOT1-containing assemblysomes. Nat Struct Mol Biol, 26(2), 110–120. 10.1038/s41594-018-0179-5

Park, S., Roelofs, J., Kim, W., et al. (2009). Hexameric assembly of the proteasomal ATPases is templated through their C termini. Nature, 459(7248), 866–870. 10.1038/nature08065

Payan-Bravo, L., Fontalva, S., Penate, X., et al. (2021). Human prefoldin modulates co-transcriptional pre-mRNA splicing. Nucleic Acids Res, 49(11), 6267–6280. 10.1093/nar/gkab446

Prakash, S., Tian, L., Ratliff, K. S., et al. (2004). An unstructured initiation site is required for efficient proteasome-mediated degradation. Nat Struct Mol Biol, 11(9), 830–837. 10.1038/nsmb814

Reincheckel, T., Sitte, N., Ullrich, O., et al. (1998). Comparative resistance of the 20S and 26S proteasome to oxidative stress. Biochem J. 10.1042/bj3350637

Reinheckel, T., Ullrich, O., Sitte, N., et al. (2000). Differential impairment of 20S and 26S proteasome activities in human hematopoietic K562 cells during oxidative stress. Arch Biochem Biophys, 377(1), 65–68. 10.1006/abbi.2000.1717

Roelofs, J., Park, S., Haas, W., et al. (2009). Chaperone-mediated pathway of proteasome regulatory particle assembly. Nature, 459(7248), 861–865. 10.1038/nature08063

Saeki, Y. (2017). Ubiquitin recognition by the proteasome. J Biochem, 161(2), 113–124. 10.1093/jb/mvw091

Saeki, Y., Toh, E. A., Kudo, T., et al. (2009). Multiple proteasome-interacting proteins assist the assembly of the yeast 19S regulatory particle. Cell, 137(5), 900–913. 10.1016/j.cell.2009.05.005

Schmidt, M., Haas, W., Crosas, B., et al. (2005). The HEAT repeat protein Blm10 regulates the yeast proteasome by capping the core particle. Nat Struct Mol Biol, 12(4), 294–303. 10.1038/nsmb914

Schmitt, M. E., Brown, T. A., & Trumpower, B. L. (1990). A rapid and simple method for preparation of RNA from Saccharomyces cerevisiae. Nucleic Acids Res, 18(10), 3091–3092. 10.1093/nar/18.10.3091

Simons, C. T., Staes, A., Rommelaere, H., et al. (2004). Selective contribution of eukaryotic prefoldin subunits to actin and tubulin binding. J Biol Chem, 279(6), 4196–4203. 10.1074/jbc.M306053200

Sorgjerd, K. M., Zako, T., Sakono, M., et al. (2013). Human prefoldin inhibits amyloid-beta (Abeta) fibrillation and contributes to formation of nontoxic Abeta aggregates. Biochemistry, 52(20), 3532–3542. 10.1021/bi301705c

Tahmaz, I., Shahmoradi Ghahe, S., Stasiak, M., et al. (2023). Prefoldin 2 contributes to mitochondrial morphology and function. BMC Biol, 21(1), 193. 10.1186/s12915-023-01695-y

Tahmaz, I., Shahmoradi Ghahe, S., & Topf, U. (2021). Prefoldin Function in Cellular Protein Homeostasis and Human Diseases. Front Cell Dev Biol, 9, 816214. 10.3389/fcell.2021.816214

Takano, M., Tashiro, E., Kitamura, A., et al. (2014). Prefoldin prevents aggregation of α-synuclein. Brain Research, 1542, 186–194. 10.1016/j.brainres.2013.10.034

Tashiro, E., Zako, T., Muto, H., et al. (2013). Prefoldin protects neuronal cells from polyglutamine toxicity by preventing aggregation formation. J Biol Chem, 288(27), 19958–19972. 10.1074/jbc.M113.477984

Thompson, D., Hakala, K., & DeMartino, G. N. (2009). Subcomplexes of PA700, the 19 S regulator of the 26 S proteasome, reveal relative roles of AAA subunits in 26 S proteasome assembly and activation and ATPase activity. J Biol Chem, 284(37), 24891–24903. 10.1074/jbc.M109.023218

Topf, U., Suppanz, I., Samluk, L., et al. (2018). Quantitative proteomics identifies redox switches for global translation modulation by mitochondrially produced reactive oxygen species. Nat Commun, 9(1), 324. 10.1038/s41467-017-02694-8

Vainberg, I. E., Lewis, S. A., Rommelaere, H., et al. (1998). Prefoldin, a chaperone that delivers unfolded proteins to cytosolic chaperonin. Cell, 93(5), 863–873. 10.1016/s0092-8674(00)81446-4

Vaquer-Alicea, J., & Diamond, M. I. (2019). Propagation of Protein Aggregation in Neurodegenerative Diseases. Annu Rev Biochem, 88, 785–810. 10.1146/annurev-biochem-061516-045049

Varshavsky, A. (2017). The Ubiquitin System, Autophagy, and Regulated Protein Degradation. Annu Rev Biochem, 86, 123–128. 10.1146/annurev-biochem-061516-044859

Verma, R., Aravind, L., Oania, R., et al. (2002). Role of Rpn11 metalloprotease in deubiquitination and degradation by the 26S proteasome. Science, 298(5593), 611–615. 10.1126/science.1075898

Verma, R., Oania, R., Graumann, J., et al. (2004). Multiubiquitin chain receptors define a layer of substrate selectivity in the ubiquitin-proteasome system. Cell, 118(1), 99–110. 10.1016/j.cell.2004.06.014

Wehmer, M., Rudack, T., Beck, F., et al. (2017). Structural insights into the functional cycle of the ATPase module of the 26S proteasome. Proc Natl Acad Sci U S A, 114(6), 1305–1310. 10.1073/pnas.1621129114

Wilmes, G. M., Bergkessel, M., Bandyopadhyay, S., et al. (2008). A genetic interaction map of RNA-processing factors reveals links between Sem1/Dss1-containing complexes and mRNA export and splicing. Mol Cell, 32(5), 735–746. 10.1016/j.molcel.2008.11.012

Wisniewski, J. R., Zougman, A., Nagaraj, N., et al. (2009). Universal sample preparation method for proteome analysis. Nat Methods, 6(5), 359–362. 10.1038/nmeth.1322

Xie, Y., & Varshavsky, A. (2001). RPN4 is a ligand, substrate, and transcriptional regulator of the 26S proteasome: a negative feedback circuit. Proc Natl Acad Sci U S A, 98(6), 3056–3061. 10.1073/pnas.071022298

Yaffe, M. P., & Schatz, G. (1984). Two nuclear mutations that block mitochondrial protein import in yeast. Proc Natl Acad Sci U S A, 81(15), 4819–4823. 10.1073/pnas.81.15.4819

Yazgili, A. S., Meul, T., Welk, V., et al. (2021). In-gel proteasome assay to determine the activity, amount, and composition of proteasome complexes from mammalian cells or tissues. STAR Protoc, 2(2), 100526. 10.1016/j.xpro.2021.100526

