## Supplementary Figures for "Identification of a non-canonical function of prefoldin subunit 5 in proteasome assembly"

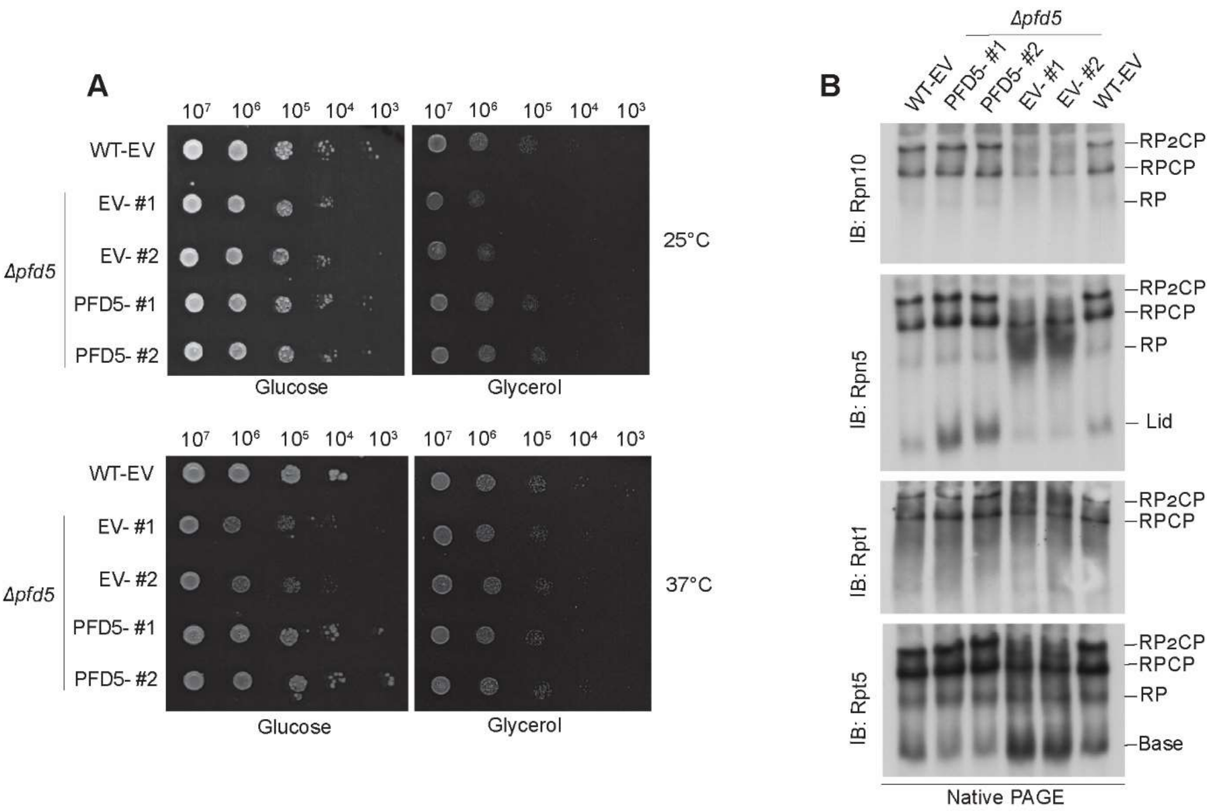


**Supplementary Fig. 1 (related to Fig. 1). Expression of *PFD5* restores the proteasome assembly in Δ*pfd5* cells. A.** The Δ*pfd5* cells were transformed with a vector expressing *PFD5* under its native promoter or empty vector (EV). Wild-type cells were also transformed with EV. Ten-fold dilutions of transformed cells were spotted on selective medium plates containing glucose or glycerol as carbon source. Cells were grown at 25°C or 37°C for 48h. Experiments were conducted in two biological repetitions. **B.** Transformed cells were grown on selective medium containing glycerol at 25°C to logarithmic growth phase. An equal concentration of cell lysates was separated by Native-PAGE and analysed by immunoblotting. WT, wild type. EV, empty vector. #1 and #2, two different yeast transformants.

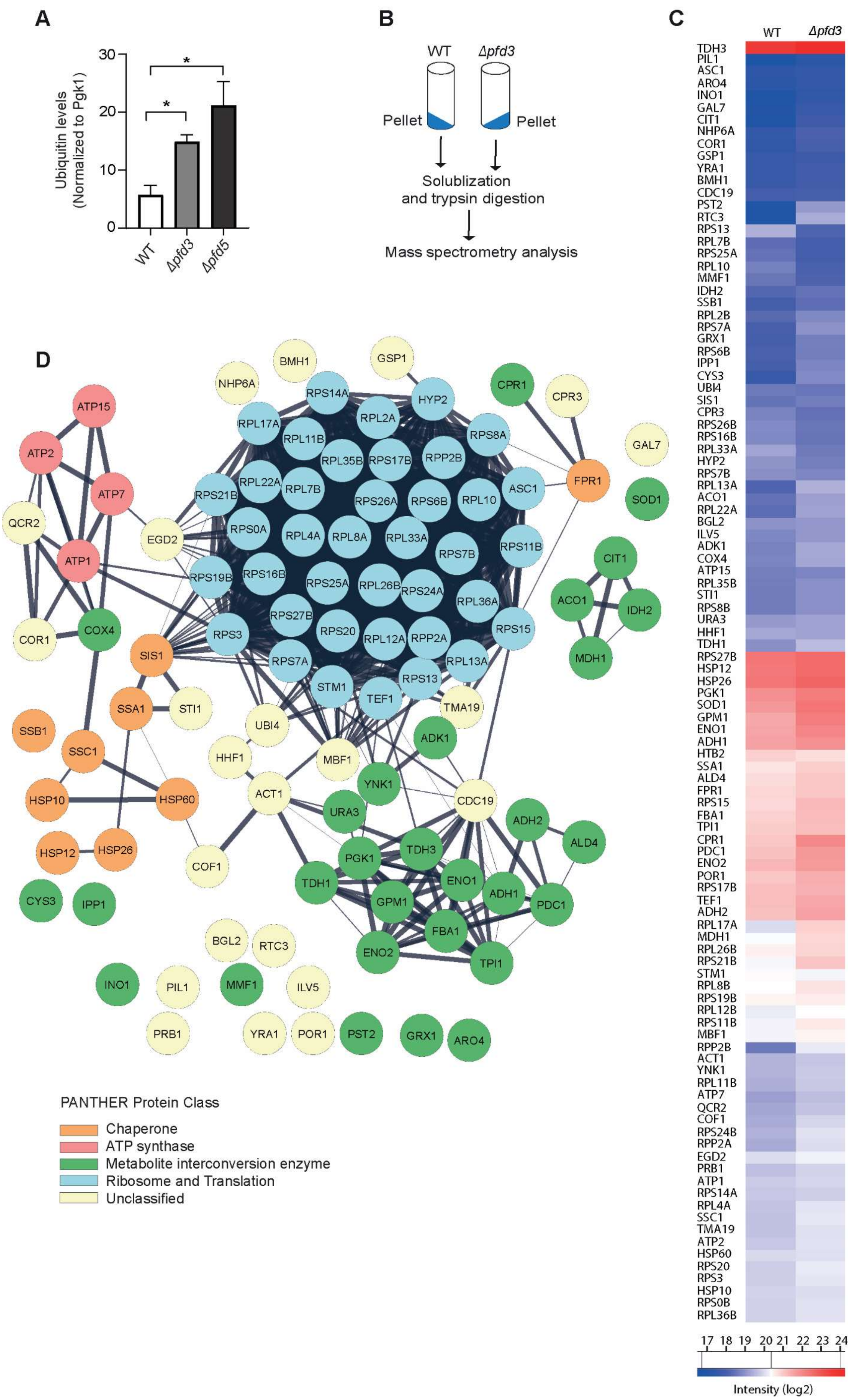
A
Ubiquitin levels (Normalized to Pgk1)
30
20
10
0
*
*
WT
Δpfd3
Δpfd5
B
WT
Δpfd3
Pellet
Pellet
Solublization and trypsin digestion
Mass spectrometry analysis
C
WT
Δpfd3
TDH3
PIL1
ASC1
ARO4
INO1
GAL7
CIT1
NHP6A
COR1
GSP1
YRA1
BMH1
CDC19
PST2
RTC3
RPS13
RPL7B
RPS25A
RPL10
MMF1
IDH2
SSB1
RPL2B
RPS7A
GRX1
RPS6B
IPP1
CYS3
UBI4
SIS1
CPR3
RPS26B
RPS16B
RPL33A
HYP2
RPS7B
RPL13A
ACO1
RPL22A
BGL2
ILV5
ADK1
COX4
ATP15
RPL35B
STI1
RPS8B
URA3
HHF1
TDH1
RPS27B
HSP12
HSP26
PGK1
SOD1
GPM1
ENO1
ADH1
HTB2
SSA1
ALD4
FPR1
RPS15
FBA1
TPI1
CPR1
PDC1
ENO2
POR1
RPS17B
TEF1
ADH2
RPL17A
MDH1
RPL26B
RPS21B
STM1
RPL8B
RPS19B
RPL12B
RPS11B
MBF1
RPP2B
ACT1
YNK1
RPL11B
ATP7
QCR2
COF1
RPS24B
RPP2A
EGD2
PRB1
ATP1
RPS14A
RPL4A
SSC1
TMA19
ATP2
HSP60
RPS20
RPS3
HSP10
RPS0B
RPL36B
17 18 19 20 21 22 23 24
Intensity (log2)
D
BMH1
GSP1
NHP6A
RPS14A
CPR1
ATP15
RPL2A
HYP2
CPR3
RPL17A
RPL11B
RPS8A
GAL7
ATP2
RPL35B
RPS17B
RPP2B
RPL22A
FPR1
ATP7
RPS21B
RPL7B
RPS26A
RPS6B
RPL10
ASC1
SOD1
QCR2
RPS0A
EGD2
RPL4A
RPL8A
RPL33A
RPS7B
RPS11B
CIT1
ATP1
RPS19B
RPS16B
RPS25A
RPL26B
RPS24A
RPL36A
ACO1
IDH2
COR1
COX4
RPS27B
RPS3
RPS20
RPL12A
RPP2A
RPS15
SIS1
RPS7A
RPL13A
MDH1
STM1
RPS13
TEF1
SSA1
STI1
TMA19
SSB1
UBI4
ADK1
SSC1
HHF1
MBF1
YNK1
ACT1
CDC19
HSP10
HSP60
URA3
ADH2
ALD4
HSP12
HSP26
PGK1
TDH3
COF1
ENO1
TDH1
ADH1
PDC1
CYS3
IPP1
GPM1
FBA1
BGL2
RTC3
ENO2
TPI1
INO1
PIL1
MMF1
ILV5
PRB1
YRA1
POR1
PST2
GRX1
ARO4
PANTHER Protein Class
Chaperone
ATP synthase
Metabolite interconversion enzyme
Ribosome and Translation
Unclassified

**Supplementary Fig. 2 (related to Fig 4). Characterisation of insoluble protein fraction in Δ*pfd3* cells A.** Quantification of ubiquitin levels in total protein extracts normalized to Pgk1. Data were presented as mean ± SEM, $n = 3$, $*p < 0.05$. **B.** Wild type and Δ*pfd3* cells were grown on full growth medium containing glycerol at 25 °C to logarithmic growth phase. Insoluble proteins were separated from cell lysate by ultracentrifugation. The pelleted proteins were solubilized, digested and analysed by mass spectrometry. **C.** Heat map comparing median LFQ intensities of four biological replicates of proteins identified in the insoluble fraction of wild-type and Δ*pfd3* cells. **D.** The interaction network of all identified insoluble proteins in Δ*pfd3* and wild type. The network was constructed with Cytoscape software (version 3.9.1) using StringApp plugin (version 1.7.1) (Doncheva et al., 2019). The colours represent the PANTHER protein classes. WT, wild type.

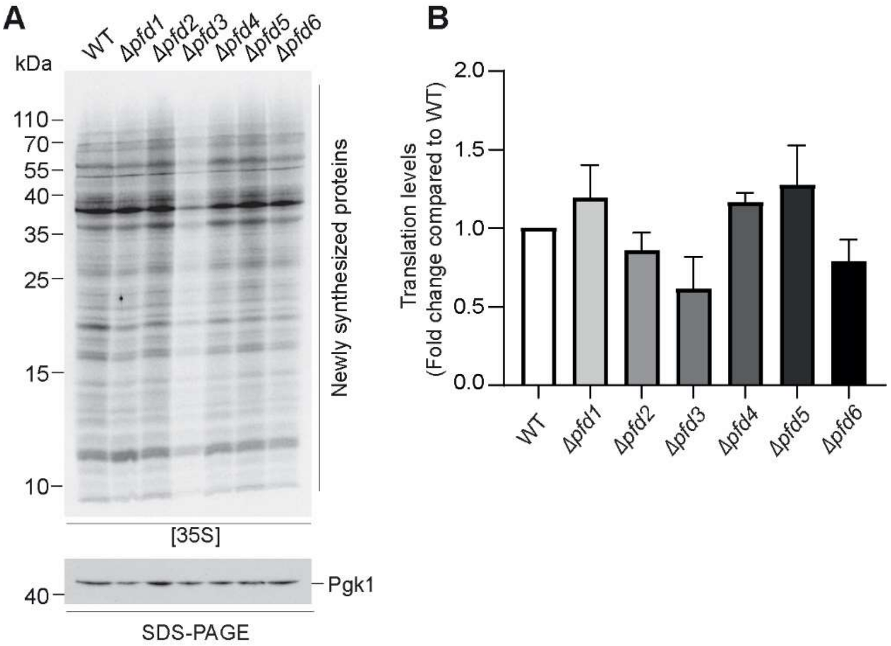


**Supplementary Fig. 3 (related to Fig. 4). Loss of prefoldin subunit 3 affects global protein synthesis. A.** Yeast cells grown to logarithmic growth phase were incubated with radiolabelled methionine. Cell lysate from equal $OD_{600}$ were loaded on SDS-PAGE and analysed by autoradiography. **B.** Quantification of translation assay shown in panel A. Autoradiography signal was normalized to Pgk1 from SDS-PAGE. Data are presented as mean ± SEM, *n* = 3.

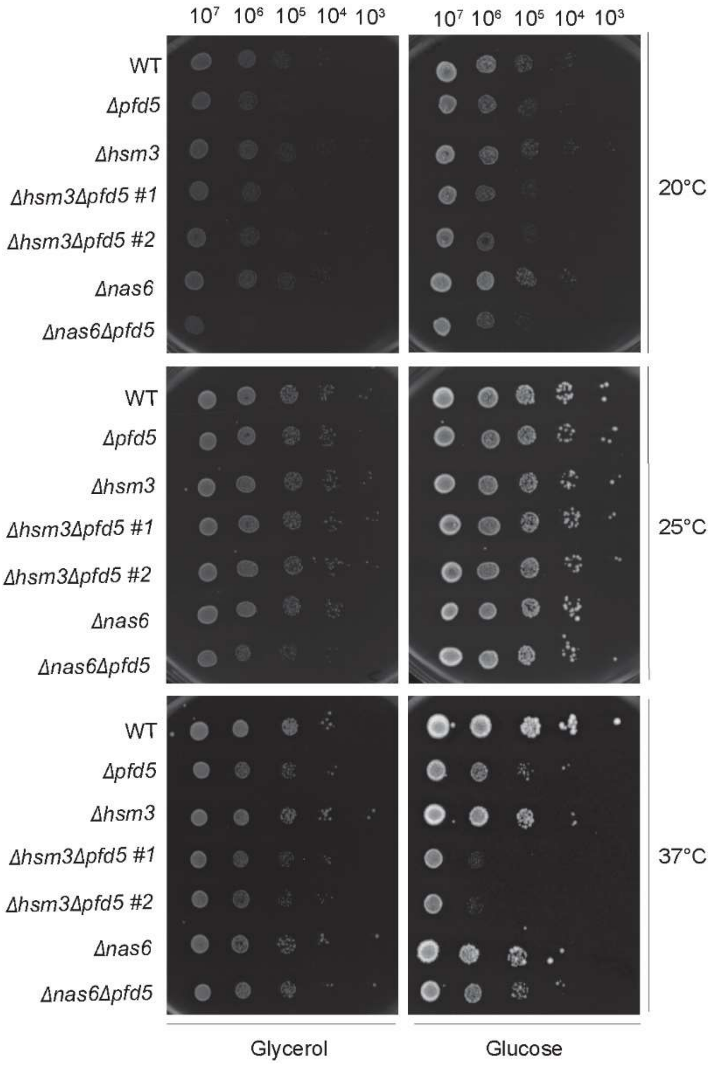


**Supplementary Fig. 4 (related to Fig. 6). Genetic interaction of *PFD5* with the Rpt-ring assembly chaperones *HSM3* and *NAS6*.** Ten-fold dilutions of wild type, single and double deletion cells were spotted on plates containing glucose or glycerol. Plates were incubated at indicated temperatures. Experiments were conducted in two biological repetitions. WT, wild type. #1 and #2, two different yeast strains obtained after generation of double deletion.

### Reference

Doncheva, N. T., Morris, J. H., Gorodkin, J., *et al.* (2019). Cytoscape StringApp: Network Analysis and Visualization of Proteomics Data. *J Proteome Res, 18*(2), 623-632. https://doi.org/10.1021/acs.jproteome.8b00702
